# Peptorama: Structure creation and exploratory data analysis for non-ribosomal peptides

**DOI:** 10.64898/2026.11.03.764117

**Authors:** Anna C. Sang, Charlotte Paton-Simpson, Theodore O. Calcott, Noah Finn, David F. Ackerley, Mark J. Calcott

## Abstract

Non-ribosomal peptides (NRPs) are a diverse family of compounds with many clinical and biotechnological uses. NRP biodiscovery increasingly focuses on biosynthetic gene clusters (BGCs), and NRP structures can generally be predicted from BGCs. Despite their chemical connection, there is no individual research tool which quickly analyses NRPs and BGCs in combination. Here, we present Peptorama, which simplifies drawing diverse NRP structures and allows rapid comparisons between known and predicted NRPs. We show the utility of peptide comparisons and interactive analysis for biodiscovery, exemplified by retrospective analysis of calcium dependent antibiotics and biodiscovery of novel pyoverdine producing candidate strains. Peptide comparisons are computationally more efficient than BGC comparisons, substantially cutting runtimes without loss of biodiscovery functionality. Peptorama can therefore enhance existing pipelines with a rapid method for comparing peptide-level diversity. It is available online at https://peptorama.com/ and is also available to be run locally.

## Introduction

Over 50% of drugs in clinical use today are derived from or inspired by naturally occurring non-ribosomal peptides (NRPs) or related NRP-polyketide hybrid products.^1^ Clinically successful NRPs have wide ranging uses, including front-line antibiotics (e.g., vancomycin, daptomycin), antifungals (e.g., caspofungin), immunosuppressants (e.g., cyclosporin A), and antitumour agents (e.g., bleomycin, dactinomycin). Their value extends beyond the clinic. Examples of this include the cyclic lipopeptide surfactin being used in bioremediation, cosmetics and crop biocontrol^2^ and pyoverdine siderophores being exploited for plant growth promotion, heavy-metal remediation and in PET imaging.^3^ Despite their success, many NRPs remain imperfect agents due to side-effects, the emergence of resistance and modest production titres. Their continued use often reflects a lack of better alternatives rather than ideal efficacy or safety, and this has driven ongoing efforts to discover new, improved analogues in nature.^4-6^

The clinical and chemical productivity of NRPs reflects how they are synthesised. NRPs are short peptides built by non-ribosomal peptide synthetase (NRPS) enzymes, multimodular assembly lines in which each module typically selects, activates and incorporates one monomer into a growing chain. This allows NRPS modules to draw from a pool of more than 500 monomer building blocks and to diversify through ribosomally inaccessible modifications such as cyclisation, oxidation, or glycosylation.^1^ It is this combinatorial flexibility that allows NRPs to expand into structural space that is not readily accessible to ribosomal synthesis or synthetic chemistry.^4^

Despite their utility, rates of NRP discovery have dropped, a major issue being that untargeted methods often rediscover known chemicals.^6,7^ Crucially, “novel chemistry” to meet a new need now seldom means a wholly new structural class. Clinically successful NRP drugs are commonly modified through synthesis and semi-synthesis to retain the validated target and mechanism while addressing the parent compound’s limitations.^1^ However, the complex structures of NRPs can make explorations of chemical space prohibitively difficult. Fortunately, evolution performs its own structure-activity optimisation to generate suites of analogues.^4-6^ Discovering these analogues in nature can overcome issues with clinically used compounds.

Several tools have been developed to guide the biodiscovery of new non-ribosomal peptides. Aiding such efforts is that the NRPS enzymes governing biosynthesis of a given NRP are often collocated in biosynthetic gene clusters (BGCs), and that candidate peptides can be predicted by analysis of the NRPS modules. For mining the unknown, antiSMASH is a predominant tool for identifying BGCs and predicting compounds,^6^ with additional tools, such as BiG-SCAPE and DisCERN, aiding the exploration of chemical diversity and discovery of new BGCs.^8,9^ Each of these tools functions by comparing BGCs in their entirety.^8,9^ While this has served as a valuable proxy, the goal is generally to identify novel compounds. A focus on comparing BGC information means that subsequent manual chemical database queries are essential to dereplicate compounds and identify leads.^6^ Current tools consequently require manual analysis and shifting between tools to identify true chemical similarity.

Here we present Peptorama, a browser-based application that allows visual comparison and exploration of known and/or predicted candidate peptides. Peptorama compares the similarity between peptides using peptide sequence identity based on monomer alignments and chemical similarity using Tanimoto similarity. It then produces a network to view alignments, related structures and associated metadata. We show that it can automate the generation of chemical structures of diverse non-ribosomal peptides, exemplified through the retrospective identification of novel calcium dependent antibiotics (CDAs), and the identification of BGCs encoding potential novel pyoverdines, for which genomic data is sparse, based on known pyoverdine structures.

## Results

### Peptorama design

Peptorama is designed to integrate chemical visualisation between known and unknown peptides. We took a user-centric approach for design, aiming to allow exploratory analysis with minimal set up. Peptorama can be installed with minimal packages and functions as a web application to allow users without command line experience to analyse data using a drag and drop user interface. The user interface has a three panelled design with the left-hand panel containing network controls and information, the central panel enabling data upload and displaying the network, and the right-hand panel displaying associated alignments, chemical structures and metadata. Small bars between each column can be clicked and dragged to adjust how much space each column occupies on the screen. This allows users, for example, to enlarge the network visualisation when they want to make a closer visual analysis. To guide users through the basic workflow, on launch, the left-most panel contains distinct orange text to attract the users’ attention to the question, “First time here?”. This panel introduces the user to the workflow as soon as possible. Example data is provided by a single click to allow users to learn about the software through direct interaction rather than relying solely on written instructions.

### Comparison of diverse peptides

A key challenge in comparing NRPs is the diversity in monomers, structures and nomenclature. To enable rapid comparisons of peptides, an automated pipeline was created to extract peptides in plain text format, identify monomers for sequence alignment and create putative structures for analysing chemical similarity (Figure 1). To allow for modified monomers to be optionally assessed as not being substantially different to their unmodified precursors, the pipeline extracts a base amino acid as well as the exact monomer sequence. The base amino acid is identified via a string comprising at least three letters in which the first character is capitalized. Characters preceding the final capitalised letter or shorter sequences separated by dashes are treated as modifications and L- and D-can optionally indicate stereotypes. As an example, this allows alignment of L-ornithine (Orn) and D-*N*^5^-formyl-*N*^5^-hydroxy-Orn (D-F-OH-Orn) under the broadest settings as they share the base of Orn. User options allow increased stringency of matching the entire monomer code and/or stereochemistry. Once monomers are extracted, pairwise sequence alignment is performed between all peptides and sequence identities calculated. In addition, candidate structures are created using inbuilt structures for known monomers or structures containing an undefined sidechain for unknown amino acids. Monomers contain placeholder numbering at the N-terminus and C-terminus to subsequently generate peptide structures using *in silico* ligation. The presence of numbering allows custom structures to be added to produce peptide structures for compounds bearing unusual monomers. Reflecting the diversity of NRPs, these custom monomers allow unique nomenclature to create a large diversity of structures. The peptides are then compared using Tanimoto similarity of Morgan fingerprints^10^ to identify similar compounds. This sequence and/or chemical similarity are then used to generate an interactive network of related peptides that can be visually explored to compare alignments, chemical structures and provided metadata.

**Figure 1.**
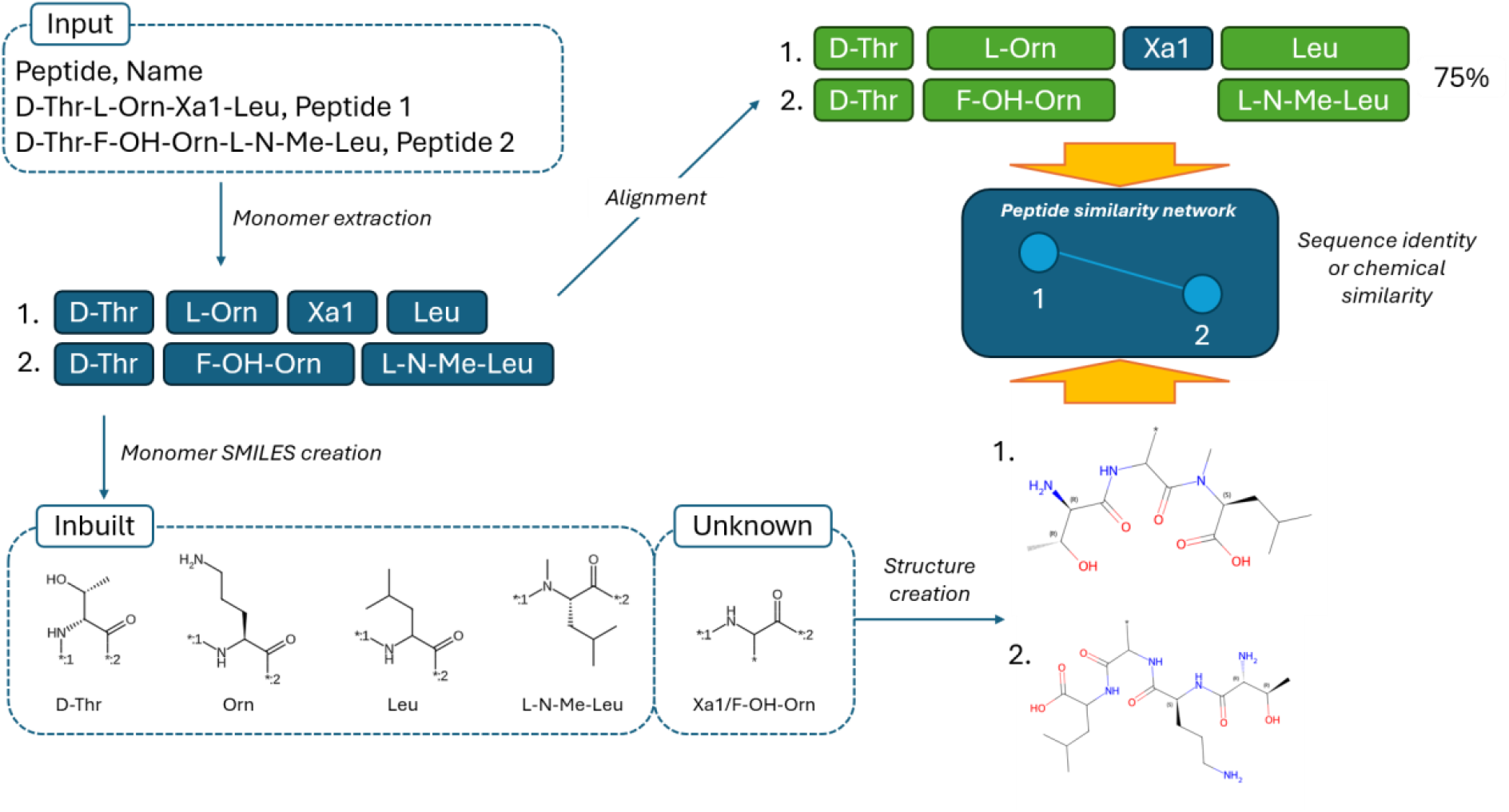
Example of processing two hypothetical peptides to create a network for comparing peptide similarity. Peptorama takes plain text or CSV formatted files as input with one peptide per line. The tool then identifies monomers and modifications based on defined but flexible rules. For sequence alignment, these monomers are aligned, with the stringency being user defined. To generate structures, monomers are generated from an inbuilt dataset when known, or a monomer with an asterisk as a sidechain when unknown. The C- and N-terminal of each monomer are indicated with placeholder values to allow *in silico* amide bond formation between adjacent monomers. The resulting structures are compared for chemical similarity based on Tanimoto similarity. An interactive network is then created in which related peptides are linked, with the option to change linkages to be based on either sequence identity or Tanimoto similarity.

### Generation of diverse structures

While structures for simple peptides are straightforward, NRPs are far more challenging due to their diversity in monomers, cross-links and cyclisations. To generate structures, chemicals are often represented as SMILES strings which are compatible with a range of software tools.^11^ The diversity of possible NRPs makes it difficult to account for every possibility, but Peptorama was designed to be capable of constructing structures from peptide strings based on the SMILES of individual monomers. While a selection of monomers is inbuilt, the placeholder numbering for the N- and C-terminus allows customized monomers to be added for any unknown monomer. To assess the flexibility of automated structure generation, we selected ten test peptides to represent a diversity of NRPs and describe key aspects below.

To consider basic cyclisation, tyrocidine A^12^ and gramicidin S^12^ were selected as head-to-tail cyclised monomeric and dimeric NRPs respectively, together with daptomycin,^13^ which is intramolecularly cyclised with the addition of N-decanoyl as the N-terminal residue (Supplementary Figure 1, Supplementary Tables 1 and 2). To represent this in peptide strings for structure creation, the cyclisation is indicated through square brackets for head-to-tail cyclisations and parentheses for intramolecular cyclisation, with multimerism introduced by additional brackets and parentheses. The N-decanoyl and non-proteinogenic residues of daptomycin were encoded using custom SMILES monomers.

**Table 1:** Benchmarking to compare runtimes and resources used with example data.

| Software | Wall time (s), mean ± SD | CPU used avg. | Peak RAM |
|---|---|---|---|
| BiG-SCAPE | 249 ± 15 | 6.7 cores | 2.00 GiB |
| DiscERN | 308 ± 1 | 4.5 cores | 1.18 GiB |
| Peptorama | 5.1 ± 0.1 | 1.3 cores | 1.67 GiB |

In addition to amide bonds, there are instances such as syringolin A and yersiniabactin in which NRPs contain other types of linkages between monomers (Figure 2a, Supplementary Tables 1 and 2). Syringolin A contains an unusual ureido linkage between two monomers instead of a peptide bond, and a malonate derivative introduced by a polyketide synthase module.^14^ The ureido linkage could be added by incorporating a custom ureidoVal monomer, defined using the base Val monomer and custom SMILES monomer to represent the unusual linkage. The malonate derivative introduces a monomer through a carbon-carbon bond. This requires the upstream amino acid to have the carbonyl removed, which is automated by introducing a forward slash between the monomers, allowing monomer ligation without having to define a different SMILES monomer for the upstream amino acid. Next, yersiniabactin is a five monomer NRP that is unusual in possessing no amide bonds despite containing three Cys residues.^15^ The Cys residues are modified by cyclisation domains that cause internal cyclisation between the thiol group of the sidechain and carbonyl of the amide bond to form thiazoline and thiazolidine rings. Automated carbonyl removal and customised monomers allow this structure to be constructed.

**Figure 2.**
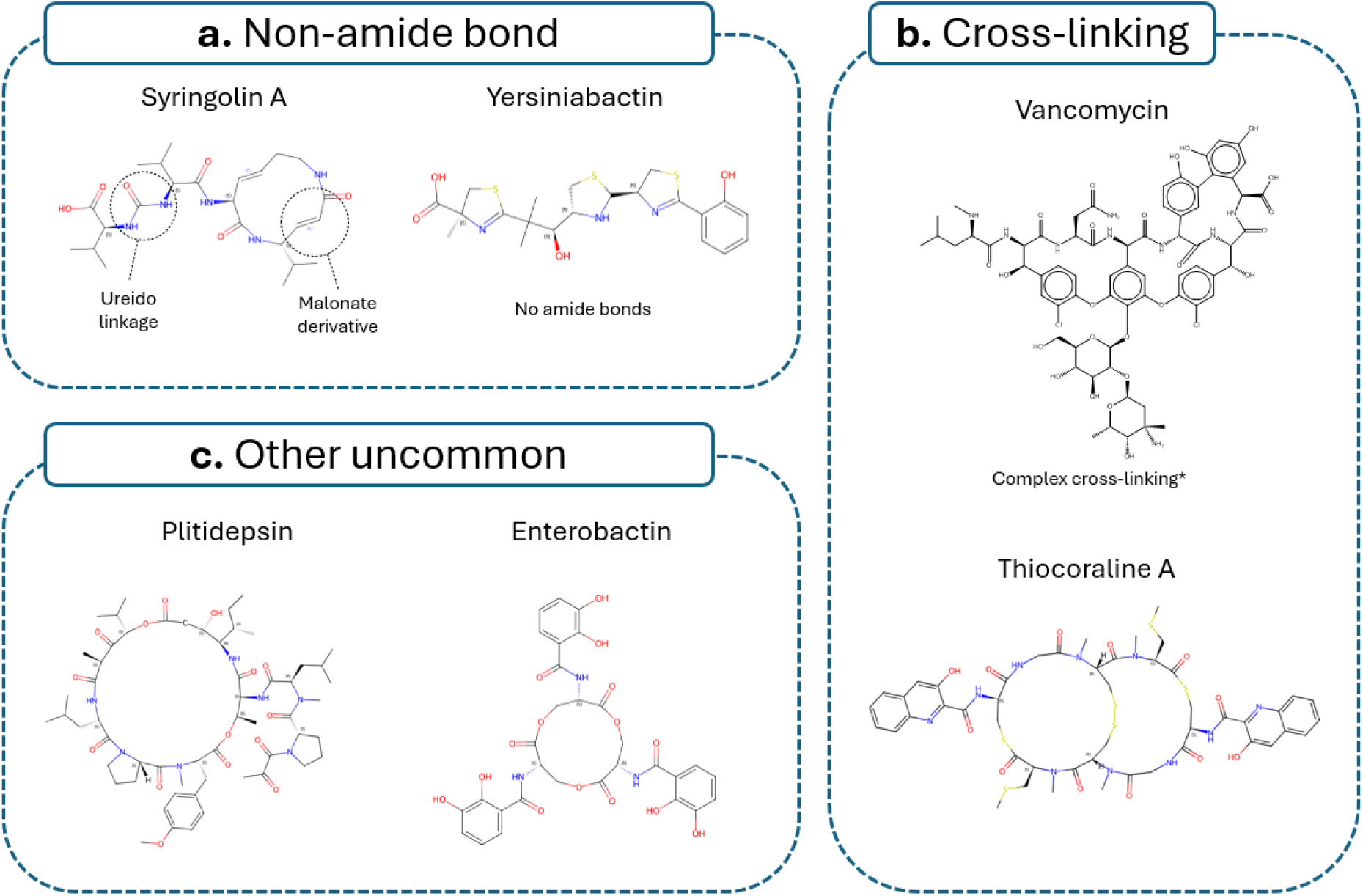
Examples of difficult-to-generate NRP structures successfully generated by Peptorama. **a.**Structures that contain bonds other than amide bonds between monomers. **b**. Structures with cross-linking between monomer sidechains. **c**. Other examples of unusual NRPs. All structures were generated in Peptorama except for vancomycin. The structure for vancomycin was considered low quality purely due to aesthetic defects (Supplementary Fig 2). The above structure of vancomycin was instead made by pasting the Peptorama generated SMILES into ChemDraw.

Other NRPs such as vancomycin and thiocoraline A exhibit cross-linking between amino acid sidechains (Figure 2b, Supplementary Tables 1 and 2). Vancomycin is a particularly difficult example to portray due to multiple cross-links and glycosylation.^16^ Peptorama created this structure using custom monomers and placeholders to indicate regions for automated cross-linking. While a valid SMILES string was created, the generated structure was visually assessed as lower quality (Supplementary Figure 2), and the complex cross-linking means that care will be required to create diverse structures bearing alternative monomers. Nonetheless, while the automated structure was lower quality (Supplementary Figure 2), this was only aesthetic with the generated SMILES being confirmed to be correct when pasted into ChemDraw (Figure 2b). Thiocoraline A is another example of cross-linking and additionally contains S-methylation, N-methylation and dimeric head-to-tail intramolecular cyclisation.^17^ The cross-linking required a custom monomer that shares a base Cys amino acid, and dimerization was indicated with double parentheses. The monomer for double methylation was represented as L-S-Me-N-Me-Cys with the N- and S-methylation being automatically added to the base Cys amino acid using an “N-Me” and “S-Me” prefix. This automated methylation allows diversity without having to define unique SMILES monomers. In contrast to vancomycin, the resulting structure was considered good quality.

Additional examples of unusual NRPs are cyclosporine A (Supplementary Figure 2, Supplementary Tables 1 and 2), plitidepsin and enterobactin (Figure 2c, Supplementary Tables 1 and 2). Cyclosporine A has an intramolecular cyclisation and was selected as an extreme example of backbone N-methylation (seven instances).^18^ Methylation was automated as per the thiocoraline A example. Plitidepsin contains N-Me-O-Me-Tyr (a double methylated Tyr analogue), isostatine and hydroxyisovalerylpropionyl (Hip).^19^ Unlike typical α-amino acids, isostatine is a β-hydroxy-γ-amino acid and Hip is an α-hydroxy acid. These required user defined SMILES monomers, and methylation was automated to generate the structure. In contrast, enterobactin is composed of three dipeptides of 2,3-dihydroxybenzoic acid and L-Ser. The three dipeptides are linked by intramolecular heterocyclisation between the C-terminus and sidechain of L-Ser residues on different dipeptides.^20^ This differs from general cyclisation between the terminal residue and sidechain of a non-terminal residue. Nonetheless, the structure was successfully generated using three parentheses as per the standard representation for intramolecular cyclisation.

### Retrospective biodiscovery of antibiotics

The discovery of malacidin highlights the value of discovering new NRPs. Malacidin is a CDA, a term describing a group of antibiotics including daptomycin, friulimicin, laspartomycin, and A54145. Malacidin is an outlier in this group, lacking a conserved motif that is found in every other known CDA.^21^ Recently, the software package DiscERN was developed to genome mine for evolutionary related natural products and discovered discomycin based on relatedness to the CDA from *Streptomyces coelicolor*.^9^ As an example of retrospective biodiscovery, to determine whether Peptorama could help identify differences between known and newly discovered compounds, we generated a network to compare malacidin and discomycin with other CDAs, to guide their discovery alongside tyrocidine A and gramicidin S as an outgroup (Figure 3a). This showed clustering of malacidin and discomycin with the other CDAs and a separate cluster for gramicidin S and tyrocidine A. Under the broadest alignment settings, allowing matches between amino acids sharing the same base such as L-Me-Asp and L-Asp, malacidin had fewest linkages to CDAs and shared 58% and 55% sequence identity with riulimicin and laspartomycin, respectively (Figure 3b). In the malacidin biodiscovery paper, an important feature of malacidin was reported to be its absence of a conserved Asp-X-Asp-Gly motif. Inspection of the sequence alignments generated in Peptorama showed this difference, however, it also identified a similar motif that had three of the four motif residues present under the broadest alignment settings (Figure 3b). This finding highlights the utility of Peptorama for discovering important relationships and visualising differences between NRPs.

**Figure 3.**
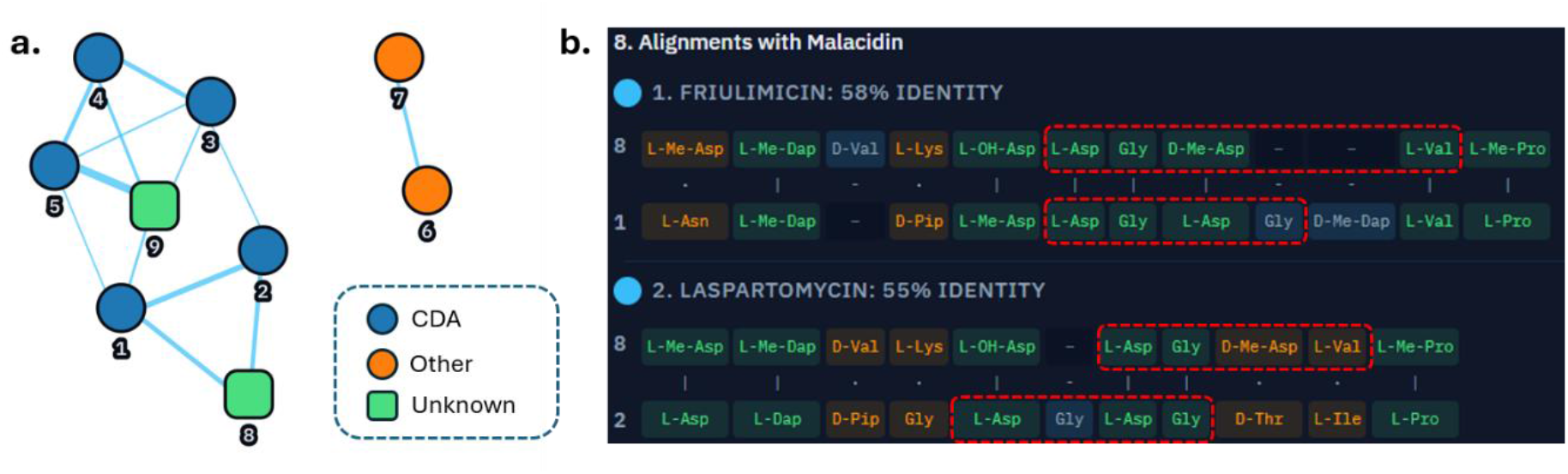
Comparing known CDAs and antibiotics to recently discovered CDAs. **a.**A network with edges representing at least 35% identity between two NRPs. **b**. Alignments between malacidin and related CDAs. The calcium binding motifs are highlighted by red dashes. NRPs are: Friulimicin; 2. Laspartomycin; 3. Daptomycin; 4. A54145; 5. CDA from *S. coelicolor*; 6. Tyrocidine; 7. Gramicidin S; 8: Malacidin; 9: Discomycin.

### Biodiscovery of pyoverdine siderophores and comparison to other tools

Pyoverdines were selected as a class of NRP to compare tools for discovering and analysing new BGCs. Pyoverdines are siderophores secreted by fluorescent pseudomonads to chelate and uptake iron. They consist of three parts, an acyl chain, a chromophore, and a peptide chain. They make good candidates for assessing biodiscovery potential, as 83 pyoverdine variants differing in their peptide chains are already known.^3^ The genomic information underpinning the biosynthesis of known pyoverdines is sparser, and we were only able to identify 22 genomes for strains that produce a known pyoverdine.

To compare to existing BGC tools, we analysed genomic predictions of pyoverdine gene clusters from known genomes and all predicted compounds from a range of newly sequenced isolated strains. For the latter, 30 strains were isolated from soil and stream water using media that is selective for fluorescent pseudomonads. The genomic DNA was isolated and sequenced for all strains. Consistent with enrichment for pseudomonads, 26 strains were confirmed as being of the *Pseudomonas* genus by comparison to a custom database using Sourmash (Supplementary Table 3). All 30 genomes plus the 22 known pyoverdine BGCs were processed with antiSMASH and analysed using BiG-SCAPE, DisCERN and Peptorama. Peptorama was the clear leader in speed and computational cost, completing the analysis in 5.1 ± 0.1 s (mean ± SD, n = 3) on a single core, with a peak memory use of 1.67 GiB. By way of comparison, BiG-SCAPE completed the benchmark in 249 ± 15 s, with an average CPU use of 6.7 cores and peak memory use of 2.00 GiB. DiscERN required 308 ± 1 s, with an average CPU use of 4.5 cores and peak memory use of 1.18 GiB. This represents a substantial increase in speed with Peptorama running 48.8X faster than the BGC-based tools.

A key advantage of Peptorama is its utility in comparing known and unknown compounds. We next used Peptorama to compare the antiSMASH predictions from 30 genomes to all 83 known pyoverdines peptide chains. The initial network had clear clustering of known pyoverdine peptide chains with a linked subcluster. There was an additional cluster with the pyoverdine chromophores and a few small clusters and individual nodes (Supplementary Figure 3). The linked subcluster was due to three pyoverdine BGCs containing two predicted products. This provided a bridge between pyoverdine peptide chain BGCs and an unrelated group of BGCs (Supplementary Figure 4). These dual NRPS clusters were manually split, and the data reanalysed to separate the linked cluster (Figure 4). All products in the peptide chain cluster, consistent with known pyoverdine peptide chains,^3^ contained two potential iron binding residues. There were seven predicted compounds in the peptide chain cluster that were consistent with encoding novel pyoverdines (Supplementary Figure 5).

**Figure 4.**
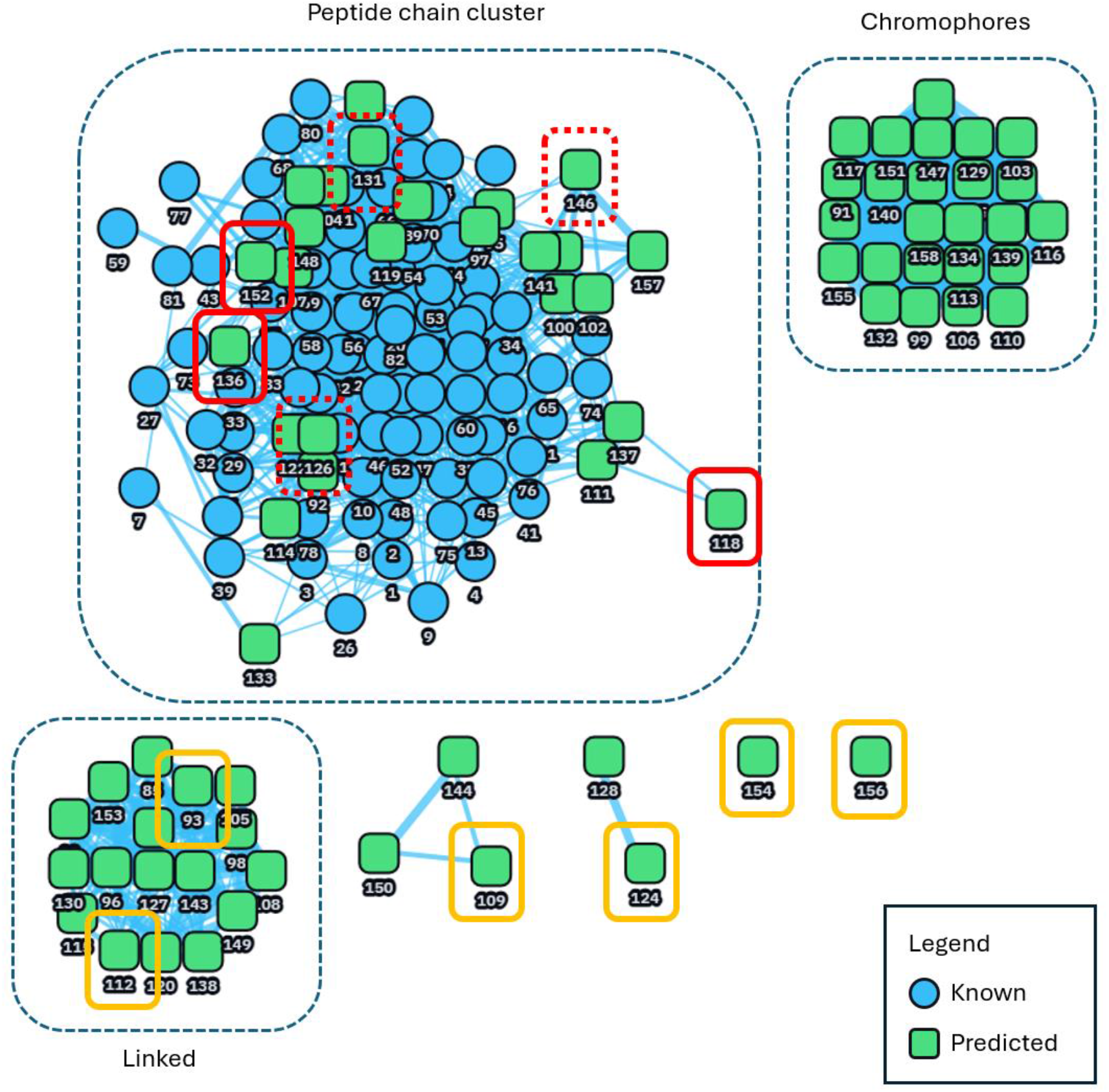
Peptorama clustering of known pyoverdines alongside antiSMASH predicted peptides from 30 isolated strains. Clustering of peptide chains, chromophores and the linked subcluster is indicated. Boxes with red solid lines indicate predicted novel pyoverdine side chains. Boxes with red dashed lines indicate gene clusters consistent with a known pyoverdine. Boxes with orange solid lines indicate example compounds outside of a pyoverdine gene cluster.

To examine the biodiscovery potential of Peptorama, as highlighted in Figure 4, three groups of compounds were selected to represent BGCs encoding three of the most novel peptide chains, three known peptide chains and six unrelated BGCs. The three nodes selected to correspond to known pyoverdine peptide chains were identical to a known compound except for node 146, which contained two unknown residues, but was the same length and matched for all defined residues (Supplementary Figure 6). The three sets of nodes were analysed using the cluster blast feature of antiSMASH. All six nodes within the peptide chain cluster contained two pyoverdines in the top three cluster blast hits, namely Pf-5 pyoverdine and pyoverdine SXM-1 (Supplementary Table 4). Node 118 also matched to azotobactin, a close pyoverdine analogue that has a modified chromophore, and which has here been defined as a pyoverdine.^3^ These are the only pyoverdines in the MiBIG database,^22^ the source for known cluster blast. No clear differentiation was observed in the antiSMASH outputs between the BGCs encoding the three most novel peptide chains and known peptide chains. In contrast, all six nodes selected outside the peptide chain cluster contained no pyoverdine predictions in the top three hits, supporting our opinion that Peptorama meaningfully clusters related BGCs and compounds.

The selected nodes from Peptorama were additionally compared using the output from BiG-SCAPE and DisCERN. Comparisons to the network generated by BiG-SCAPE showed the nodes for these compounds were dispersed, with no indication to distinguish pyoverdines or novelty (Supplementary Figure 7). DisCERN identified compounds within the pyoverdine peptide chain cluster as having higher K values, i.e. more likely to be pyoverdine-like BGCs, however, the most novel pyoverdines had lower K values and there was overlap in the range of K values between all three groups of compounds (Supplementary Table 5). Consequently, while relatedness to known compounds and BGCs is highlighted in Peptorama, both BiG-SCAPE and DisCERN require further manual inspection of gene clusters to confirm relatedness.

## Discussion

Several tools have been developed to identify novel BGCs, however, the underlying reason for searching for novelty is to identify new compounds. Using current tools, researchers must subsequently rely on manual searches to find novel compounds. Peptorama has been developed to automate both structure creation and searching for new NRPs. The structure creation enables broad representation of a range of NRPs, allowing for cyclisation, cross-linking and monomers to all be customised. We show the potential for Peptorama to be used for biodiscovery through retrospective analysis of CDAs and analysis of putative pyoverdines from *Pseudomonas* isolates. Peptorama clustered CDAs and highlighted differences between commonly known CDAs and recent discoveries using sequence alignment. It further showed that malacidin bears a related motif to the calcium binding region, which was reported as being absent in its discovery.^21^ When analysing *Pseudomonas* genomes for potentially novel pyoverdines, Peptorama showed a cluster containing all known pyoverdine peptide chains. Analysis of this cluster showed BGCs in this cluster were most similar to known pyoverdine BGCs, and nodes outside this region were predicted to be most similar to unrelated BGCs. This clear separation of other nodes was not present using other tools. Overall, we show Peptorama to be a valuable complement to existing biodiscovery tools, using peptide similarity to rapidly compare known and predicted structures.

## Methods

### Identifying pyoverdine gene clusters

Genomes for pyoverdine producing strains were identified from the original report or based on searches for the related strain in the NCBI database. In instances in which only Illumina data was available, draft genomes were assembled de novo using SPAdes v4.3.0. Assemblies were polished using BWA-MEM2 v2.2.1, SAMtools v1.24 and Pilon v1.24. Assembly quality and completeness were assessed using QUAST v.5.3.0 and CheckM2 v1.1.0, respectively. Final assemblies were annotated using Prokka v.1.15.6. Where only gene sequences were found for a strain in the NCBI database (typically gene sequences used for classification), the genome for the closest related strain was identified. Genome GenBank files were processed with antiSMASH and the results were analysed to find a BGC that matched the peptide chain of the known pyoverdine. The GenBank file for this BGC was subsequently rerun through antiSMASH again to generate a reference antiSMASH output for the pyoverdine BGC.

### Isolating environmental bacteria

Soil and stream water samples were collected from four separate locations around Wellington, New Zealand (ōtari-Wilton’s Bush Reserve, Khandallah Park, Seton Nossiter Park and Trelissick Park). Two samples were collected from both soil and stream water at three collection sites. Each site was a separate location along the stream. Strains were isolated by adapting a previously established method and dilutions plated on Gould’s S1 medium agar plates containing 100 µg/ml cycloheximide.^23^ Plates were incubated for three days in the dark at room temperature. Colonies were subsequently streaked on LB agar plates and incubated for two days at RT. Individual colonies were inoculated into 3 ml of LB and incubated at 18 °C, 200 rpm. Sterile glycerol was then added to 40% (v/v) and stocks were stored at −80 °C.

A total of 30 isolates had genomes sequenced by PlasmidSaurus. This included an isolate from each duplicate for ōtari-Wilton’s Bush Reserve and a single isolate from each site for the other three locations. Isolates were grown overnight in LB at 18 °C, 200 rpm. Samples were prepped and resuspended in Zymo 1X DNA/RNA Shield™ as per PlasmidSaurus cell preparation details. Genomes were sequenced by PlasmidSaurus using Oxford Nanopore Technologies (ONT) long-read sequencing. De novo assembly and species identification (via Sourmash) were performed using the Plasmidsaurus bioinformatics pipeline. The resulting GenBank files were processed using antiSMASH.

### Media

Gould’s S1 medium contained (per litre): 18 g agar, 10 g sucrose, 10 ml of glycerol, 5 g of Casamino acids, 1 g NaHCO_3_, 1 g MgSO_4_ · 7H_2_O, 2.3 g K_2_HPO_4_, 1.2 g sodium lauroyl sarcosine and 100 µg/ml mg of trimethyloprim. Trimethyloprim and sucrose was added once the media was cool and the final pH was 7.4 to 7.6.^24^

### Bench marking

BiG-SCAPE (v2.0.3), DiscERN (v.0.1.0) and Peptorama (v0.1.0) were each run three times on the antiSMASH output of the 30 test genomes (424 regions) and the 22 reference BGCs for known pyoverdines, using a workstation with an Intel Core i5-10400 CPU (8 logical processors available to the analyses) and 7.6 GiB RAM under WSL2. BiG-SCAPE and DiscERNs wall-clock time, CPU utilisation and peak memory were recorded with GNU time (/usr/bin/time -v). Peptorama was timed through its web interface, run in headless Firefox (v157.0.1). CPU time for Peptorama was the CPU time of the Firefox process tree plus the Peptorama server, and peak memory was the highest summed resident memory of the Firefox processes plus the server’s peak. Installation, database and model downloads, Pfam preparation, and browser and server start-up were excluded. Results are mean ± SD of three runs. The benchmarking dataset is available from: https://doi.org/10.5281/zenodo.23897768

## Code availability

Source code for Peptorama is available from: https://github.com/MarkCalcott/peptorama.

## Acknowledgements

This work was supported by a Health Research Council of New Zealand Hercus Fellowship awarded to M.J.C. (contract 23/188/A).

## Contributions

M.C. conceived the work, with additional ideas contributed by A.S., C.P. and D.A. Strain collection and isolation was performed by M.C. and T.C. Data analysis and image preparation was completed by M.C. and A.S. Benchmarking was performed by A.S. Software development was completed by M.C. and design was completed by C.P. and M.C. Software testing was performed by A.S., N.F., C. P. and M.C. The first draft of the manuscript was by A.S, C.P., T.C. D.A and M.C, and all authors approved the final version.

## References

1 Agrawal, S., Acharya, D., Adholeya, A., Barrow, C. J. & Deshmukh, S. K. Nonribosomal Peptides from Marine Microbes and Their Antimicrobial and Anticancer Potential. Front Pharmacol 8, 828 (2017). 10.3389/fphar.2017.00828

2 Qiao, J. et al. Research advances in the identification of regulatory mechanisms of surfactin production by Bacillus: a review. Microbial Cell Factories 23, 100 (2024). 10.1186/s12934-024-02372-7

3 Calcott, M. & Ackerley, D. in Preprints (Preprints, 2026).

4 Calcott, M. J., Sang, A. C., Ackerley, D. F. & Little, R. F. Evolutionary insights and guidelines to achieve effective and high-yield non-ribosomal peptide and polyketide engineering. Current Opinion in Microbiology 91, 102755 (2026). 10.1016/j.mib.2026.102755

5 Albarano, L., Esposito, R., Ruocco, N. & Costantini, M. Genome Mining as New Challenge in Natural Products Discovery. Marine Drugs 18, 199 (2020).

6 Hemmerling, F. & Piel, J. Strategies to access biosynthetic novelty in bacterial genomes for drug discovery. Nature Reviews Drug Discovery 21, 359–378 (2022). 10.1038/s41573-022-00414-6

7 Yang, J. Y. et al. Molecular Networking as a Dereplication Strategy. J Nat Prod 76, 1686–1699 (2013). 10.1021/np400413s

8 Draisma, A. et al. BiG-SCAPE 2.0 and BiG-SLiCE 2.0: scalable, accurate and interactive sequence clustering of metabolic gene clusters. Nat Commun 17, 2000 (2026). 10.1038/s41467-026-68733-5

9 Owen, J. G., Woolly, E. F., Lai, H.-E., Woolner, V. H. & Little, R. F. DiscERN: an automated genome mining tool for the discovery of evolutionarily related natural products. Nat Commun 17, 8657 (2026). 10.1038/s41467-026-75491-x

10 Rogers, D. & Hahn, M. Extended-Connectivity Fingerprints. Journal of Chemical Information and Modeling 50, 742–754 (2010). 10.1021/ci100050t

11 Nguyen-Vo, T.-H., Teesdale-Spittle, P., Harvey, J. E. & Nguyen, B. P. Molecular representations in bio-cheminformatics. Memetic Computing 16, 519–536 (2024). 10.1007/s12293-024-00414-6

12 Hotchkiss, R. D. The Chemical Nature of Gramicdin and Tyrocidine. Journal of Biological Chemistry 141, 171–185 (1941).

13 Micklefield, J. Daptomycin Structure and Mechanism of Action Revealed. Chem Biol 11, 887–888 (2004). 10.1016/j.chembiol.2004.07.001

14 Clerc, J. et al. Synthetic and structural studies on syringolin A and B reveal critical determinants of selectivity and potency of proteasome inhibition. Proceedings of the National Academy of Sciences 106, 6507–6512 (2009). 10.1073/pnas.0901982106

15 Drechsel, H. R. et al. Structure elucidation of yersiniabactin, a siderophore from highly virulent Yersinia strains. Liebigs Annalen 1995, 1727–1733 (1995).

16 Sheldrick, G. M., Jones, P. G., Kennard, O., Williams, D. H. & Smith, G. A. Structure of vancomycin and its complex with acetyl-D-alanyl-D-alanine. Nature 271, 223–225 (1978). 10.1038/271223a0

17 Perez Baz, J., Cañedo, L. M., Fernández Puentes, J. L. & Silva Elipe, M. V. Thiocoraline, a novel depsipeptide with antitumor activity produced by a marine Micromonospora. II. Physico-chemical properties and structure determination. J Antibiot (Tokyo) 50, 738–741 (1997). 10.7164/antibiotics.50.738

18 Mikol, V., Kallen, J., Pflügl, G. & Walkinshaw, M. D. X-ray structure of a monomeric cyclophilin A-cyclosporin A crystal complex at 2.1 A resolution. J Mol Biol 234, 1119–1130 (1993). 10.1006/jmbi.1993.1664

19 Alonso-Álvarez, S. et al. Plitidepsin: design, development, and potential place in therapy. Drug Design, Development and Therapy 11, 253–264 (2017). 10.2147/DDDT.S94165

20 Raymond, K. N., Dertz, E. A. & Kim, S. S. Enterobactin: An archetype for microbial iron transport. Proceedings of the National Academy of Sciences 100, 3584–3588 (2003). doi:10.1073/pnas.0630018100

21 Hover, B. M. et al. Culture-independent discovery of the malacidins as calcium-dependent antibiotics with activity against multidrug-resistant Gram-positive pathogens. Nat Microbiol 3, 415–422 (2018). 10.1038/s41564-018-0110-1

22 Zdouc, Mitja M. et al. MIBiG 4.0: advancing biosynthetic gene cluster curation through global collaboration. Nucleic Acids Res 53, D678–D690 (2025). 10.1093/nar/gkae1115

23 Butaitė, E., Baumgartner, M., Wyder, S. & Kümmerli, R. Siderophore cheating and cheating resistance shape competition for iron in soil and freshwater Pseudomonas communities. Nat Commun 8, 414 (2017). 10.1038/s41467-017-00509-4

24 Gould, W. D., Hagedorn, C., Bardinelli, T. R. & Zablotowicz, R. M. New selective media for enumeration and recovery of fluorescent pseudomonads from various habitats. Appl Environ Microbiol 49, 28–32 (1985). 10.1128/aem.49.1.28-32.1985

