## Supplementary material for "Peptorama: Structure creation and exploratory data analysis for non-ribosomal peptides"

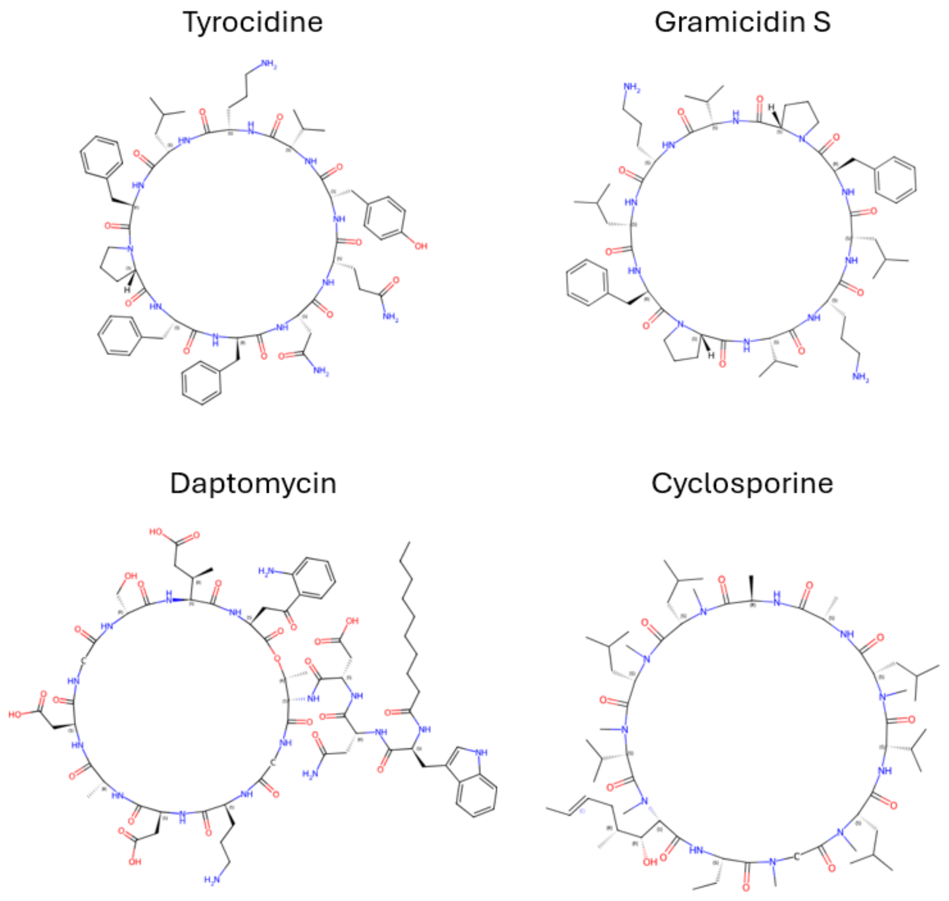


Figure 1. Examples of peptide structures generated in Peptorama.

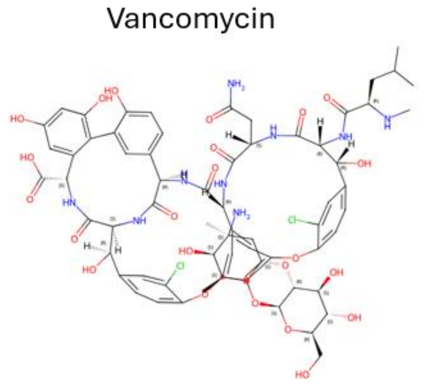


Figure 2. Peptorama generated structure of Vancomycin. The SMILES string was confirmed to be correct, however, the peptide did not display well due to complex cyclisation.

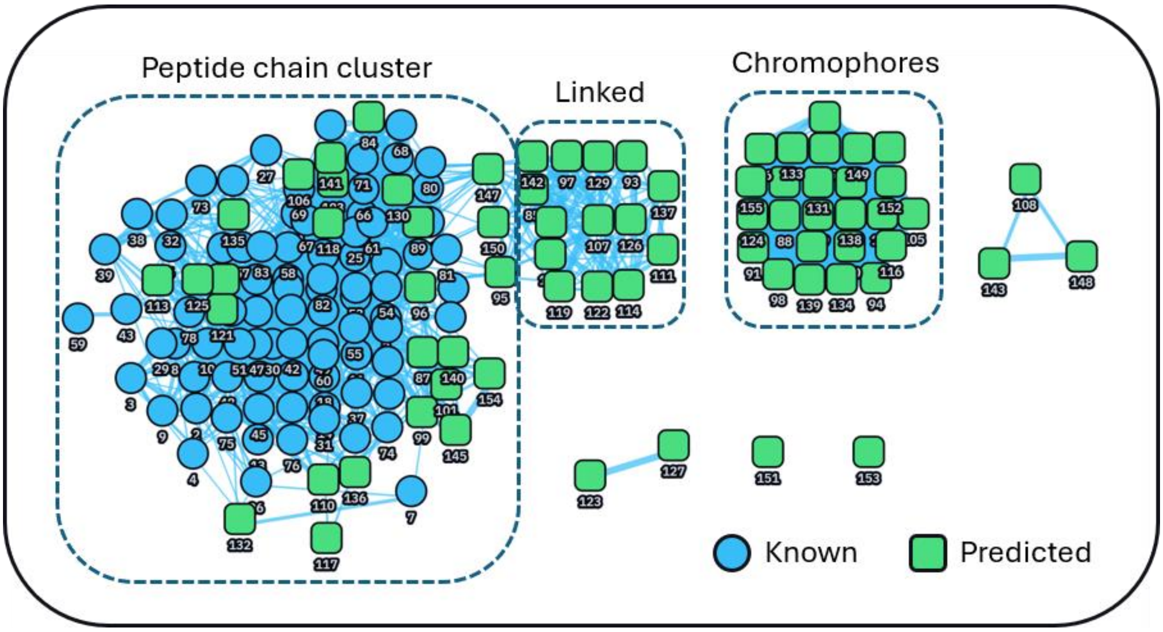


Figure 3. Network generated in Peptorama prior to manually splitting the linked subcluster. Known pyoverdine peptide chains are represented by blue circles, and antiSMASH predicted peptides from 30 genomes of isolated bacteria are represented as green squares.

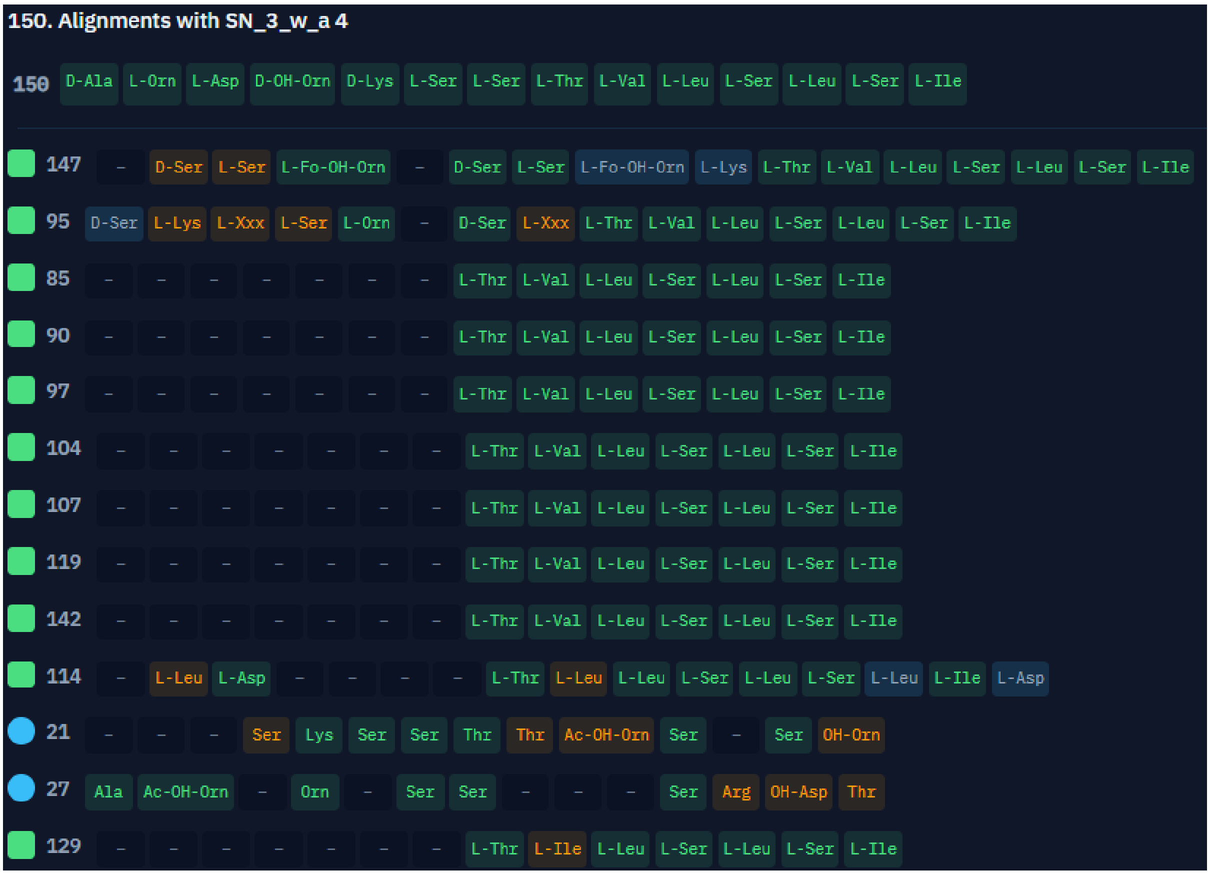


Figure 4. Compact alignment representation showing alignments to peptide 150. Peptides 95, 147 and 150 from the bridge between the sidechain cluster and linked cluster. Peptides 85, 90, 97, 104, 104, 107, 114, 119, 129 and 142 align primarily with the C-terminal region of peptide 150. Numbering based on input into Peptorama. Blue circles indicate a match to a known pyoverdine peptide chains and green squares to a predicted peptide.

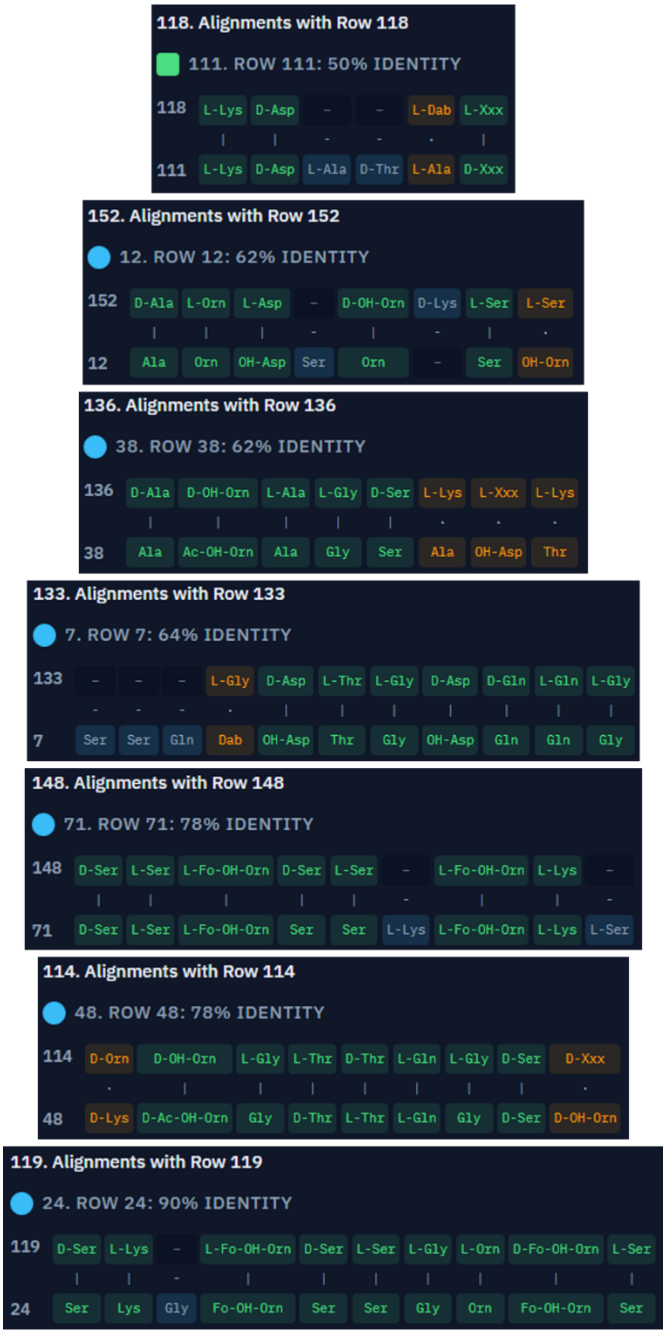


*Figure 5. Predicted peptides differing from known pyoverdines with the closest alignment to another peptide.*

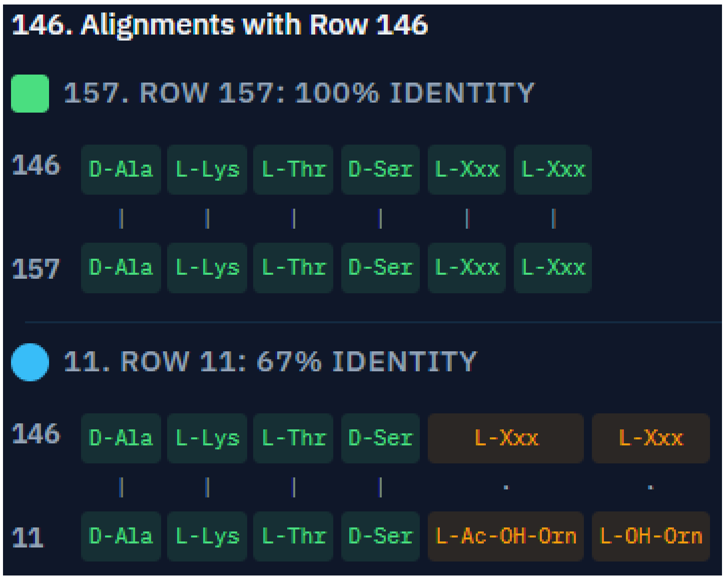


*Figure 6. Alignment showing node 146 differing from node 11, i.e. the closest known compound, in not having two residues defined.*

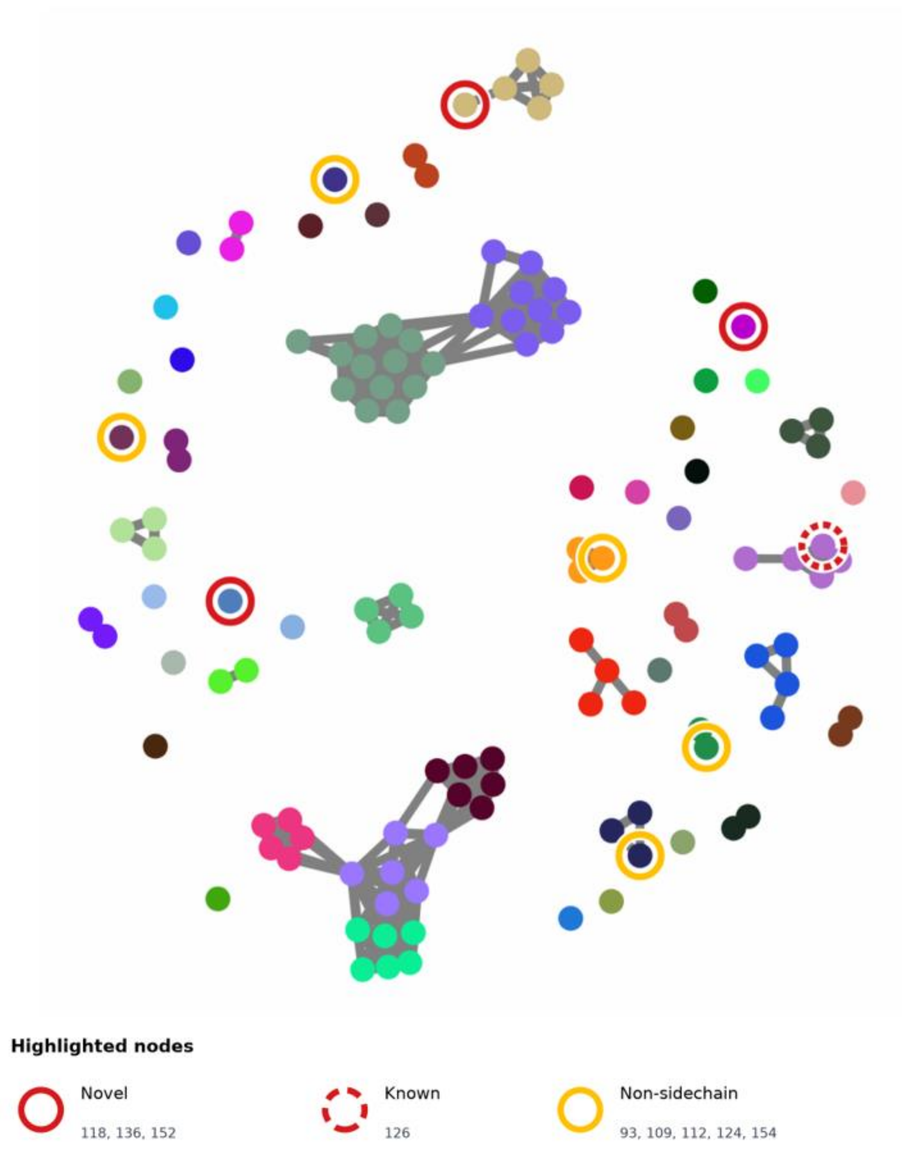


Figure 7. BiG-SCAPE clustering of known pyoverdines with antiSMASH predictions from 30 isolated strains. Circles with red solid lines indicate BGCs encoding predicted novel pyoverdine side chains. Circles with red dashed lines indicate BGCs consistent with a known pyoverdine. Circles with orange solid lines indicate example BGCs outside of a pyoverdine gene cluster. Nodes 131, 146, and 156 are not shown because their regions are hybrid clusters (NRPS with hydrogen-cyanide, ranthipeptide, or betalactone) which BiG-SCAPE placed in a different bin to the reference containing NRPS bin. Node 131 and 156 were placed in NRPS.other and node 146 was plcaed in NRPS.RiPP.

Table 1. Peptide strings used to draw structures for select non-ribosomal peptides.

| Name | Peptide |
|---|---|
| Tyrocidine A | [D-Phe-L-Pro-L-Phe-D-Phe-L-Asn-L-Gln-L-Tyr-L-Val-L-Orn-L-Leu] |
| Gramicidin S | [[D-Phe-L-Pro-L-Val-L-Orn-L-Leu]] |
| Daptomycin | n-Dec-L-Trp-D-Asn-L-Asp-(L-Thr-Gly-L-Orn-L-Asp-D-Ala-L-Asp-Gly-D-Ser-L-3-Me-Glu-L-Kyn) |
| Syringolin A | L-ureidoVal-L-Val-(L-dhLys-L-Val/DH-Mal) |
| Yersiniabactin | Sal/L-Thz/L-Thd/di-Me-KR-Mal/L-Me-Thz |
| Vancomycin | D-N-Me-Leu-D-Bht-L-Asn-D-gvHpg-D-Hpg-L-Bht-L-Dpg |
| Thiocoraline A | 3HQA-((D-Cys-Gly-L-N-Me-cl-Cys-L-S-Me-N-Me-Cys)) |
| Cyclosporine | [D-Ala-L-N-Me-Leu-L-N-Me-Leu-L-N-Me-Val-L-N-Me-Bmt-L-Abu-N-Me-Gly-L-N-Me-Leu-L-Val-L-N-Me-Leu-L-Ala] |
| Plitidepsin | Pyr-L-Pro-D-N-Me-Leu-(L-Thr-isoSta-Hip-L-Leu-L-Pro-L-N-Me-O-Me-Tyr) |
| Enterobactin | DHB-(((L-Ser))) |

Table 2. Custom monomers used to draw structures for select non-ribosomal peptides.

| name | code | smiles |
|---|---|---|
| beta-hydroxytyrosine | Bht | [*:1]NC([C@H](O)C1=CC(Cl)=C(O([*:5]))C=C1)C(=O)[*:2] |
| (4R)-4-[(E)-2-butenyl]-4-methyl-l-threonine | Bmt | [*:1]NC([C@H](O)[C@H](C)C\\C=C\\C)C(=O)[*:2] |
| N-decanoyl | n-Dec | CCCCCCCCCC(=O)[*:2] |
| 2-3-dihydroxybenzoic acid | DHB | C1(C(O)=C(O)C=CC=1)C(=O)[*:2] |
| 3-5-dihydroxyphenylglycine | Dpg | [*:1]NC(C1=C([*:6])C(O)=CC(O)=C1)C(=O)[*:2] |
| 3-Me-Glu | 3-Me-Glu | [*:1]NC([C@H](C)CC(=O)O[*:3])C(=O)[*:2] |
| Hydroxyisovalerylpropionyl | Hip | [*:1]O[C@@H](C(C)C)C(=O)[C@H](C)C(=O)[*:2] |
| 4-hydroxyphenylglycine | Hpg | [*:1]NC(C1=CC([*:6])=C(O)C=C1)C(=O)[*:2] |
| Glucose + vancosamine 4-hydroxyphenylglycine | gvHpg | [*:1]NC(C1=CC([*:5])=C(O[C@@H]2O[C@H](CO)[C@@H](O)[C@H](O)[C@@H](O[C@@H]3O[C@@H](C)[C@@H](O)[C@@](C)(N)C3)2)C([*:5])=C1)C(=O)[*:2] |
| 3-hydroxyquinaldic acid | 3HQA | C1(O)=CC2=C(C=CC=C2)N=C1C(=O)[*:2] |
| Kynurenine | Kyn | [*:1]NC(CC(=O)C1=C(N)C=CC=C1)C(=O)[*:2] |
| 3-4-dehydrolysine | dhLys | [*:1]NC(/C=C/CCN[*:3])C(=O)[*:2] |
| Dimethylated / reduced malonate | di-Me-KR-Mal | [*:1][C@@H](O)C(C)(C)C(=O)[*:2] |
| Pyruvic acid | Pyr | [*:1]CC(=O)C(=O)[*:2] |
| Salicylate | Sal | C1C=CC=C(O)C=1C(=O)[*:2] |
| Isostatine | isoSta | [*:1]N[C@H]([C@@H](C)CC)[C@@H](O)CC(=O)[*:2] |
| Thiazolidine | Thd | [*:1]C1SC[C@H](N1)C(=O)[*:2] |
| Thiazoline (methylated) | L-Me-Thz | [*:1]C1SC[C@@](C)(N=1)C(=O)[*:2] |
| Thiazoline | L-Thz | [*:1]C1SC[C@@H](N=1)C(=O)[*:2] |
| Uriedo-linked Val | L-ureidoVal | [*:1]C(=O)(O)[C@H](C(C)C)NC(=O)[*:2] |
| Cross-linked cysteine | cl-Cys | [*:1]NC(CS([*:5]))C(=O)[*:2] |

Table 3. Species and location of isolated strains.

| Sample name | Species | Location | Source |
|---|---|---|---|
| KP_1_d_a | *Pseudomonas arsenicoxydans* | Khandallah Park | Dirt |
| KP_1_w_a | *Pseudomonas sp003852405* | Khandallah Park | Stream |
| KP_2_d_a | *Pseudomonas laurylsulfatiphila* | Khandallah Park | Dirt |
| KP_2_w_b | *Pseudomonas sp003852405* | Khandallah Park | Stream |
| KP_3_d_a | *Pseudomonas allokribbensis* | Khandallah Park | Dirt |
| KP_3_w_a | *Pseudomonas cyclaminis* | Khandallah Park | Stream |
| OW_1_d_a | *Pseudomonas fluorescens CD* | Ōtari-Wilton's Bush Reserve | Dirt |
| OW_1_d_b | *Pseudomonas fluorescens CD* | Ōtari-Wilton's Bush Reserve | Dirt |
| OW_1_w_a | *Pseudomonas lurida* | Ōtari-Wilton's Bush Reserve | Stream |
| OW_1_w_b | *Pseudomonas canadensis* | Ōtari-Wilton's Bush Reserve | Stream |
| OW_2_d_a | *Pseudomonas sp016651305* | Ōtari-Wilton's Bush Reserve | Dirt |
| OW_2_d_b | *Pseudomonas koreensis A* | Ōtari-Wilton's Bush Reserve | Dirt |
| OW_2_w_a | *Pseudomonas sp037994325* | Ōtari-Wilton's Bush Reserve | Stream |
| OW_2_w_b | *Pseudomonas viridiflava D* | Ōtari-Wilton's Bush Reserve | Stream |
| OW_3_d_a | *Pseudomonas allokribbensis* | Ōtari-Wilton's Bush Reserve | Dirt |
| OW_3_d_b | *Pseudomonas allokribbensis* | Ōtari-Wilton's Bush Reserve | Dirt |
| OW_3_w_a | *Pseudomonas lactis* | Ōtari-Wilton's Bush Reserve | Stream |
| OW_3_w_b | *Pseudomonas sp943914515* | Ōtari-Wilton's Bush Reserve | Stream |
| SN_1_d_a | *Pseudomonas sp001655615* | Seton Nossiter Park | Dirt |
| SN_1_w_a | *Pseudomonas neuropathica* | Seton Nossiter Park | Stream |
| SN_2_d_a | *Pseudomonas fluorescens CC* | Seton Nossiter Park | Dirt |
| SN_2_w_b | *Pseudomonas sp013385985* | Seton Nossiter Park | Stream |
| SN_3_d_a | *Pseudomonas putida_AE* | Seton Nossiter Park | Dirt |
| SN_3_w_a | *Pseudomonas cyclaminis* | Seton Nossiter Park | Stream |
| TP_1_d_a | *Klebsiella grimontii* | Trelissick Park | Dirt |
| TP_1_w_a | *Pseudomonas bubulae* | Trelissick Park | Stream |
| TP_2_d_a | *Acinetobacter bohemicus* | Trelissick Park | Dirt |
| TP_2_w_b | *Klebsiella terrigena* | Trelissick Park | Stream |
| TP_3_d_a | *Klebsiella ornithinolytica* | Trelissick Park | Dirt |
| TP_3_w_a | *Pseudomonas sp036884845* | Trelissick Park | Stream |

*Table 4. Top three hits from known cluster blast using antiSMASH.*

| node | strain | contig | region | type | KnownClusterBlast1 | KnownClusterBlast2 | KnownClusterBlast3 |
|---|---|---|---|---|---|---|---|
| 118 | OW_2_w_a | 1 | 5 | novel | Pf-5 pyoverdine | azotobactin D | pyoverdine SXM-1 |
| 136 | SN_1_d_a | 1 | 15 | novel | Pf-5 pyoverdine | pyoverdine SXM-1 | viscosin |
| 152 | SN_3_w_a | 2 | 4 | novel | viscosin | pyoverdine SXM-1 | Pf-5 pyoverdine |
| 126 | OW_3_d_b | 1 | 5 | known | Pf-5 pyoverdine | pyoverdine SXM-1 | viscosin |
| 131 | OW_3_w_a | 1 | 5 | known | Pf-5 pyoverdine | pyoverdine SXM-1 | viscosin |
| 146 | SN_3_d_a | 1 | 8 | known | Pf-5 pyoverdine | pyoverdine SXM-1 | viscosin |
| 93 | KP_3_d_a | 1 | 7 | non-sidechain | lokisin | anikasin | bananamide 1/bananamide 2/bananamide 3 |
| 109 | OW_1_w_b | 1 | 6 | non-sidechain | pyochelin | enantio-pyochelin | skyllamycin D/skyllamycin E |
| 112 | OW_2_d_a | 1 | 7 | non-sidechain | anikasin | lokisin | viscosinamide A/pseudodesmin A |
| 124 | OW_3_d_a | 1 | 8 | non-sidechain | chitinimide B/chitinimide F/chitinimide D/chitinimide A/chitinimide C/chitinimide E/chitinimide G | detoxin N2/detoxin N3 | pseudovibriamide A1/pseudovibriamide B1/pseudovibriamide A2/pseudovibriamide A3/pseudovibriamide A4/pseudovibria |
| 154 | SN_3_w_a | 2 | 10 | non-sidechain | cepaciachelin | ornibactin C8/ornibactin C4/ornibactin C6 | ornibactin |
| 156 | TP_3_w_a | 1 | 6 | non-sidechain | fengycin | plipastatin | mycosubtilin |

*Table 5: K Values from DisCERN for selected nodes.*

| Node | K value | Type |
|---|---|---|
| 118 | 2 | Novel |
| 136 | 3 | Novel |
| 152 | 0 | Novel |
| 126 | 3 | Known |
| 131 | 3 | Known |
| 146 | 3 | Known |
| 93 | 0 | Non-sidechain |
| 109 | 0 | Non-sidechain |
| 112 | 0 | Non-sidechain |
| 124 | 0 | Non-sidechain |
| 154 | 0 | Non-sidechain |
| 156 | 0 | Non-sidechain |

*Node numbering as per Table 4.*
